# Cyclin B1 scaffolds MAD1 at the corona to activate the spindle assembly checkpoint

**DOI:** 10.1101/726224

**Authors:** Lindsey A Allan, Magda Reis, Yahui Liu, Pim Huis in ’t Veld, Geert JPL Kops, Andrea Musacchio, Adrian T Saurin

## Abstract

The Cyclin B:CDK1 kinase complex is the master regulator of mitosis that phosphorylates hundreds of proteins to coordinate mitotic progression. We show here that, in addition to these kinase functions, Cyclin B also scaffolds a localised signalling pathway to help preserve genome stability. Cyclin B1 localises to an expanded region of the outer kinetochore, known as the corona, where it scaffolds the spindle assembly checkpoint (SAC) machinery by binding directly to MAD1. In vitro reconstitutions map the key binding interface to a few acidic residues in the N-terminus of MAD1, and point mutations in this region remove corona MAD1 and weaken the SAC. Therefore, Cyclin B1 is the long-sought-after scaffold that links MAD1 to the corona and this specific pool of MAD1 is needed to generate a robust SAC response. Robustness, in this context, arises because Cyclin B1-MAD1 localisation becomes MPS1-independent after the corona has been established. We demonstrate that this allows corona-MAD1 to persist at kinetochores when MPS1 activity falls, ensuring that it can still be phosphorylated on a key C-terminal catalytic site by MPS1. Therefore, this study explains how corona MAD1 generates a robust SAC signal and why stripping of this pool by dynein is essential for SAC silencing. It also reveals that the key mitotic kinase, Cyclin B1-Cdk1, scaffolds the pathway that inhibits its own degradation.

## INTRODUCTION

During mitosis all duplicated chromosomes must attach correctly to microtubules so they can segregate properly when the cell divides. This attachment is mediated via the kinetochore, which is a giant molecular complex assembled on chromosomes at the centromere [1]. As well as attaching to microtubules, the kinetochore must also regulate this process to ensure it occurs correctly. One aspect of this regulation involves the activation of the spindle assembly checkpoint (SAC), which blocks mitotic exit until all kinetochores have attached to microtubules. The principle of the SAC is that each unattached kinetochore acts as a factory to produce an inhibitor of mitotic exit, known as the mitotic checkpoint complex or MCC (for further molecular details see [2]). The generation of MCC is so efficient that every single kinetochore signalling centre must eventually be extinguished by microtubule attachment to allow the cell to exit mitosis [3, 4].

This complicated inactivation process, known as SAC silencing, requires the removal of catalysts that are needed at unattached kinetochores to generate the MCC [5]. Two key catalysts in this regard are MAD1, which drives the first step in MCC assembly, and MPS1, the kinase responsible for recruiting and phosphorylating MAD1 as well as other components needed for MCC assembly. Kinetochore-microtubule attachment extinguishes these activities because microtubules displace MPS1 from its binding site on NDC80 [6, 7] and at the same time they provide a highway onto which dynein motors can travel to strip MAD1 away from kinetochores [8–11]. Removal of both MPS1 and MAD1 is essential for SAC silencing because if either one is artificially tethered to kinetochores then the SAC fails to switch off and mitotic exit is blocked [12, 13].

One key unexplained aspect of the SAC concerns the kinetochore binding sites for MAD1. MAD1 is recruited to kinetochores via an established KNL1-BUB1 pathway and, in human cells, by an additional pathway involving the ROD/ZW10/Zwilch (RZZ) complex at the kinetochore’s corona (a fibrous crescent that forms around kinetochores to aid the capture of microtubules) [14]. How exactly MAD1 is recruited to the corona and whether this pool of MAD1 can signal to the SAC is unknown. It is crucial to resolve these issues because it is ultimately the RZZ complex that is stripped by dynein to shut down the SAC, implying that this pool of MAD1 is important for MCC generation [8–11]. However, the corona is positioned some distance away from MPS1 and the proposed catalytic centre for MCC generation at the KNL1/MIS12/NDC80 (KMN) network. Therefore, it remains unclear how MAD1 could signal to the SAC from the corona and it is difficult to resolve this issue without knowledge of how MAD1 binds to this region.

We show here that the key mitotic regulator – Cyclin B1/CDK1 – acts as the physical adaptor that links MAD1 to the corona. This unanticipated scaffolding function of Cyclin B1 is crucial for a robust SAC response, which we demonstrate is due the ability of corona-tethered MAD1 to respond to low level of kinetochore MPS1 activity. This study therefore reveals how the corona pool of MAD1 signals to the SAC and it explains why MPS1 inhibition and dynein-mediated stripping of the corona are both essential for SAC silencing.

## RESULTS

### Cyclin B1-MAD1 interaction facilitates Cyclin B1 and MAD1 localisation to unattached kinetochores

The Cyclin B:CDK1 kinase complex is a master regulator of mitosis that is activated during G2 phase of the cell cycle to initiate mitotic entry and degraded after chromosome alignment to induce mitotic exit. Analysis of endogenously-tagged Cyclin B1-EYFP localisation in RPE1 cells suggested that its localisation was specifically regulated during mitosis. In particular, Cyclin B1 positive foci appeared after nuclear envelope breakdown and disappeared as mitosis progressed (**Figure 1A and Movie S1**). Immunofluorescence analysis demonstrated that this localisation pattern reflects specific binding to unattached kinetochores, which is reminiscent of the checkpoint protein MAD1 (**Figure 1B-C**). In particular, Cyclin B depends on MPS1 activity to be established at this location, but thereafter it became largely insensitive to MPS1 inhibition (**Figure 1D-E**), as also shown previously for MAD1 [15]. To probe for MAD1 and Cyclin B1 association in cells, we recruited LacI-MAD1 to a LacO array on chromosome 1 in U2OS cells. This was sufficient to co-recruit Cyclin B1 in a manner that was dependent on a region between amino acids 41-92 of MAD1 (**Figure 1F-G**). Therefore, these data are consistent with earlier reports that Cyclin B1 localises to unattached kinetochores [16, 17] in a manner that is dependent on the N-terminus of MAD1 [18, 19]}.

**Figure 1.**
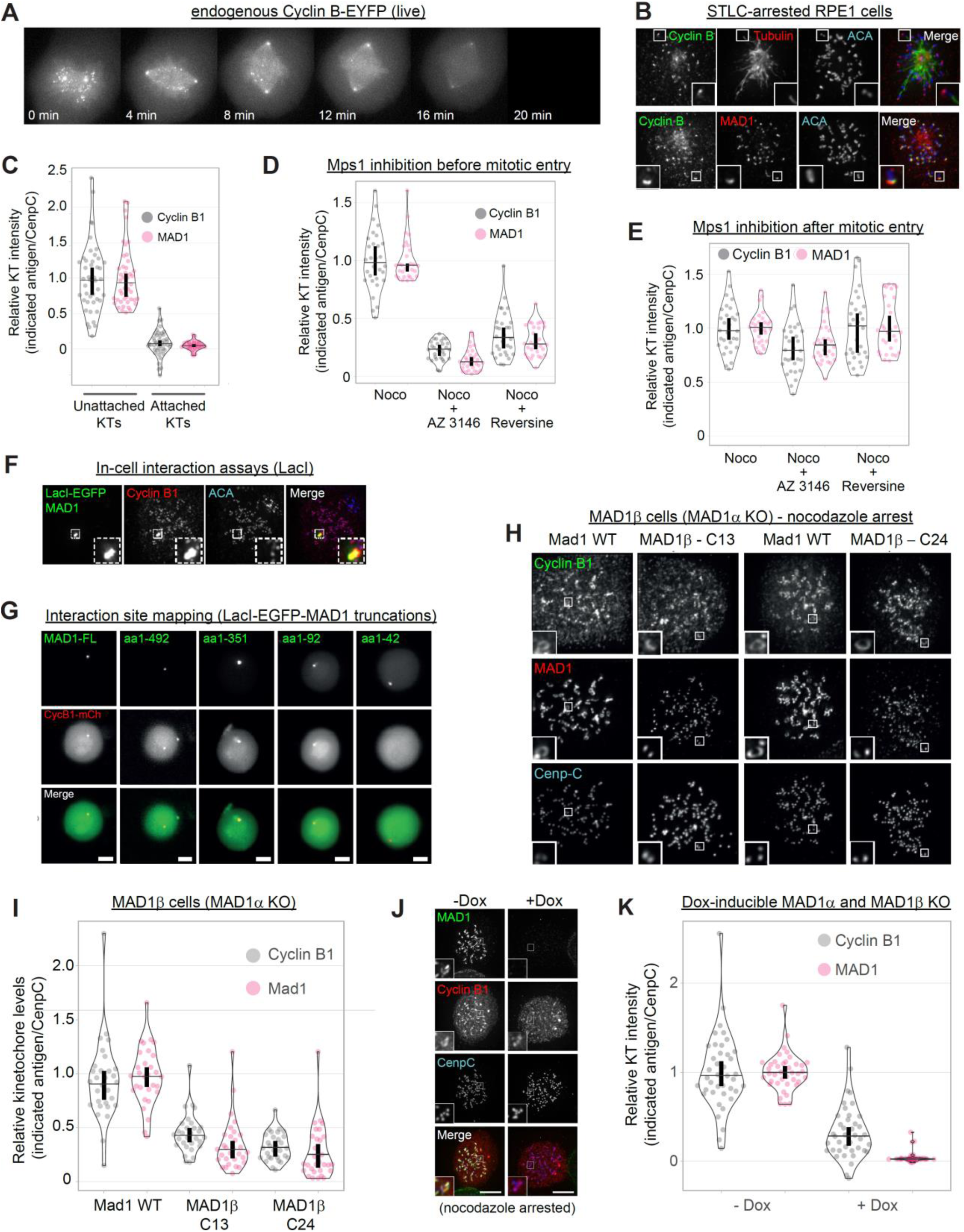
Cyclin B1-MAD1 interaction helps both proteins to localise to unattached kinetochores. **A.** Endogenous Cyclin B-YFP localisation during mitosis live in RPE 1cells. **B**,**C.** Immunofluorescence images (B) and quantifications (C) of Cyclin B and MAD1 levels at unattached and attached kinetochores in cells arrested in STLC. Each dot represents a kinetochore, data is from 40 kinetochores. Vertical bars show the 95% confidence interval. **D**,**E.** Quantification of kinetochore intensities of Cyclin B1 and MAD1 in nocodazole-arrest cells (noco) treated with the MPS1 inhibitors, AZ-3146 (5μM) or Reversine (500nM), either before (D) or after (E) mitotic entry. Each dot represents a cell and the errors bars display the variation between the experimental repeats (displayed as ±SD of the experimental means). 30 cells from 3 experiments. **F.** Immunofluorescence images of LacI-MAD1 and Cyclin B1 in U2OS cells containing a LacO arrays on chromosome 1. **G.** Live imaging of Cyclin B1-mCherry (CycB1-mCh) in LacO-U2OS cells transfected with LacI-MAD1-FL (full length: aa1-718) or various LacI-Mad1 truncations (amino acid numbers indicated). **H**,**I.** Immunofluorescence images (H) and quantifications (I) of Cyclin B and MAD1 kinetochore levels in control (Mad1-WT) or Mad1β HeLa cells (two independent clones: C13 or C24) treated with nocodazole. **J**,**K.** Immunofluorescence images (J) and quantification (K) of Cyclin B and MAD1 kinetochore localisation in doxycycline-inducible MAD1α and β knockouts treated with or without dox for 10 days and then arrested in nocodazole. Cells were selected that had full MAD1 knockout in the doxycycline treatment (this constituted approximately 30% of cells). For kinetochore intensity graphs in panel D, E, I and K each dot represents a cell, horizontal lines indicate the median and vertical bars show the 95% confidence interval. D, E and I show 30 cells from 3 experiments and K shows 40 cells from 4 experiments.

To determine the function of Cyclin B1 at kinetochores, we attempted to remove it from this location by knocking down endogenous MAD1 and replacing it with a Cyclin B1-binding defective mutant. However, all of the siRNAs tested only mildly reduced MAD1 protein (results not shown). This may be due to the fact that MAD1 is a very stable protein in cells because it takes over a week to fully deplete MAD1 following genetic deletion; see [20]). Therefore, to attempt to fully remove Cyclin B1 from kinetochores, we generated a MAD1α knockout cell line that retains only a MAD1β splice variant lacking exon 4 which encodes the Cyclin B1 binding region (hereafter referred to as MAD1β cells; **Figure S2**) [21]. Surprisingly, Cyclin B1 was reduced but still present at unattached kinetochores in MAD1β cells (**Figure 1H-I**). This was not due to residual interaction with MAD1β because doxycycline-inducible knockout of both MAD1α and MAD1β [22] completely removed MAD1 from unattached kinetochores but did not further reduce kinetochore Cyclin B1 (**Figure J-K;** note, the data shown is from 10 days doxycycline treatment which is the minimum time it takes to fully deplete endogenous MAD1 in this system). Therefore, in contrast to a recent report [18], these data demonstrate that MAD1 contributes to Cyclin B1 kinetochore localisation, but it is not the only binding partner for Cyclin B1 at kinetochores. At least one other receptor exists that is sufficient to maintain substantial levels of Cyclin B1 on kinetochores in the absence of MAD1.

Although inhibiting MAD1-Cyclin B1 interaction did not abolish Cyclin B1 recruitment to kinetochores, it did cause a dramatic effect on MAD1 localisation itself. As discussed earlier, MAD1 localises to the kinetochores via two separate pathways in human cells: the KNL1-BUB1 pathway at the outer kinetochore and the RZZ pathway at the corona. **Figure 1H** shows that Cyclin B and MAD1 both bind to the expanded corona in wild type cells, but strikingly, when their interaction is prevented in MAD1β cells, it is MAD1, and not Cyclin B1, that is lost from the corona. Therefore, this suggested that Cyclin B1 may act as a scaffold to recruit MAD1 to this region. Although MAD1 is well-known to bind the corona [14, 23–31], an interaction with a corona component has never been mapped in vitro. In fact, the only established way to remove MAD1 from the corona is to deplete RZZ subunits, which simply abolishes corona formation altogether. Therefore, we next sought to explore whether Cyclin B1 might be the receptor that directly recruits MAD1 to the corona.

### Cyclin B1 directly scaffolds MAD1 at the corona

We first assayed for direct MAD1 and Cyclin B1 interaction by obtaining homogeneously purified recombinant MAD1:MAD2 and Cyclin B1:CDK1 complexes and testing their interaction by size-exclusion chromatography (SEC), which separates proteins based on size and shape. When combined stoichiometrically with MBP-MAD1:MAD2, GST-CDK1:Cyclin B1 underwent a prominent shift in elution volume and co-eluted with MAD1:MAD2, indicative of a binding interaction (**Figure 2A;** MBP and GST are affinity tags that stand for maltose binding protein and glutathione S-transferase, respectively). Early elution of MAD1:MAD2 from the SEC column reflects its high hydrodynamic radius, typical of highly elongated structures rich in coiled-coil. As expected, the elution volume of MAD1:MAD2 was not affected by the interaction with GST-CDK1:Cyclin B1.

**Figure 2.**
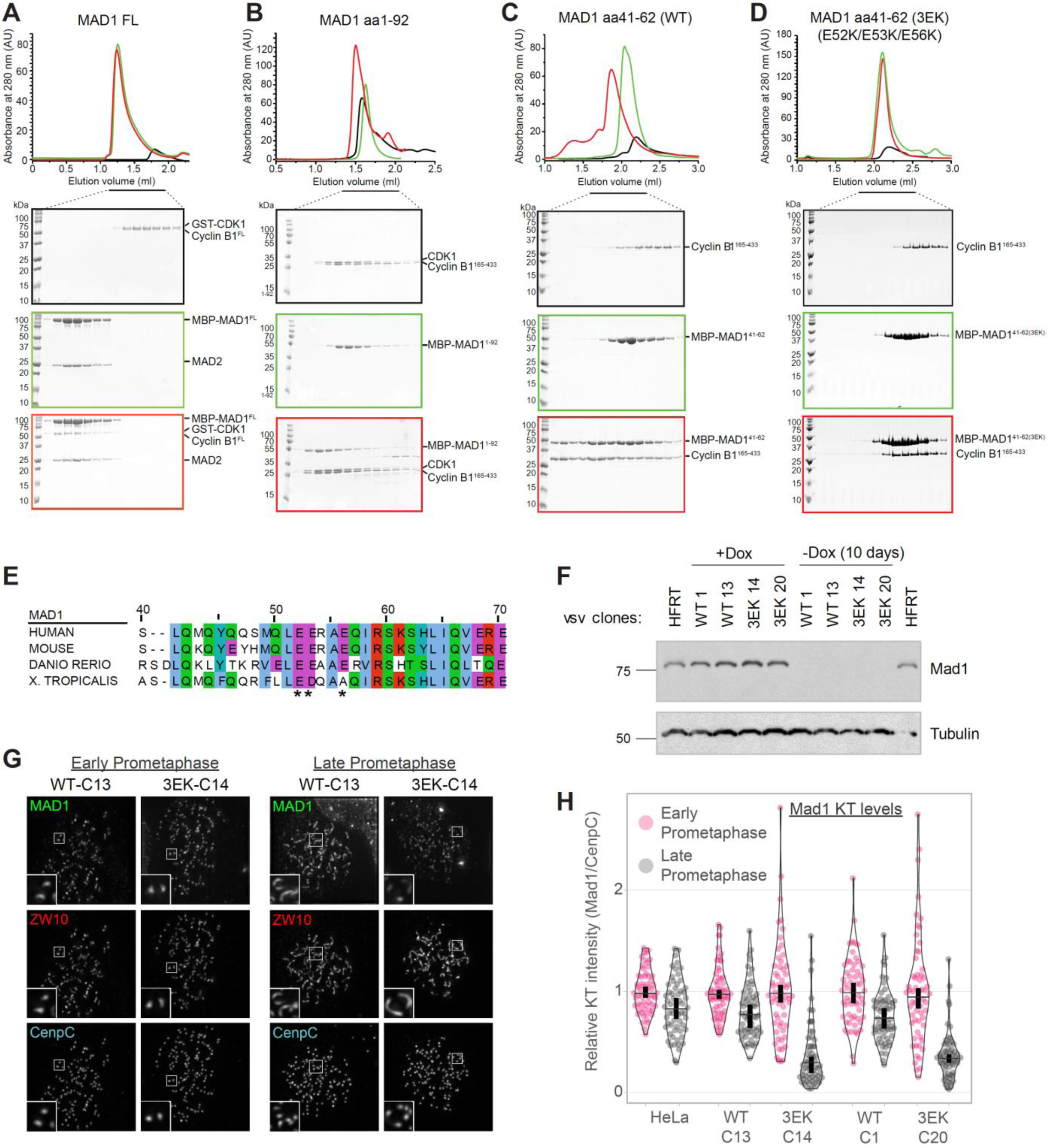
Direct Cyclin B1-MAD1 interaction is needed to localise MAD1 the corona. **A.** Elution profiles and SDS-PAGE for SEC runs on a Superose 6 Increase 5/150 GL column of the GST-CDK1:Cyclin B1 complex (black profile), MBP-MAD1:MAD2 (green profile), and their combination (red profile). FL = full length. B. Elution profiles and SDS-PAGE for SEC runs on a Superdex 200 Increase 3.2/300 column of GST-CDK1:Cyclin B1^165-433^ complex (black), MBP-MAD1^1-92^ (green), and their combination (red). **C.** Elution profiles and SDS-PAGE for SEC runs on a Superose 6 Increase 5/150 GL column of Cyclin B1^165-433^ (black), MBP-MAD1^41-62^ (green), and their combination (red). **D.** Elution profiles and SDS-PAGE for SEC runs on a Superose 6 Increase 5/150 GL column of Cyclin B1^165-433^ (black), MBP-MAD141-62 carrying the E52K, E53K, and E56K mutations (green), and their combination (red). **E.** Alignment of the N-terminal region of Cyclin B1 that contains the MAD1 binding region. Numbering refers to the HsMAD1 sequence. **F.** Western blot analysis of indicated vsv-MAD1-WT or 3EK HeLa clones treated with or without doxycycline for 10 days. **G.** Immunofluorescence images showing MAD1 and ZW10 kinetochore levels in nocodazole-arrested MAD1-WT-C13 AND 3EK-C14 just after nuclear envelope breakdown (early prometaphase) or later in mitosis when the chromatin is condensed (late prometaphase). **H.** Quantification of MAD1 kinetochore localisation from indicated MAD1-WT and 3EK clones treated as in (G). Each dot represents a cell, horizontal lines indicate the median and vertical bars show the 95% confidence interval. 60 cells from 3 experiments.

In the absence of Cyclin B1, GST-CDK1 did not interact directly with MAD1:MAD2 (**Figure S3A**). Cyclin B1, on the other hand, interacted with MAD1:MAD2, albeit more weakly than was observed with the GST-CDK1:Cyclin B1 complex (**Figure S3B**). Removal of residues 1-92 from MAD1 (MAD1^93-718^:MAD2) abolished the interaction with Cyclin B1 (**Figure S3C**), indicating that residues 1-92 of MAD1 are necessary for the interaction. Importantly, the N-terminal region of MAD1 was also sufficient to bind Cyclin B1:CDK1, as revealed by SEC experiments with MBP-MAD1^1-92^ pand CDK1:Cyclin B1^165-433^ (**Figure 2B**; note, the Cyclin B1 N-terminal region was removed in the experiments to improve yield, stability, and solubility). Therefore, MAD1 binds directly to Cyclin B1:CDK1 via a region located in the first 92 residues of MAD1.

To narrow this region down further, we performed additional truncations of MAD1^1-92^, which led to the identification of a minimal Cyclin B1:CDK1 binding site in residues 41-62 of MAD1 (**Figure S3D-F;** the full list of tested constructs is in **Table S1, Part I**). Importantly, MAD1^41-62^ interacted not only with CDK1:Cyclin B1_165-433_, but also with the isolated Cyclin B1^165-433^, thus confirming results in **Figure S3** that Cyclin B1 contains the MAD1 binding site (**Figure 2C and S3F**). To identify determinants of the MAD1^41-62^:Cyclin B1 interaction, we extensively mutagenized residues in the MAD1^41-62^ segment and on Cyclin B1 (as summarized in **Table S1, Parts II and III**; selected examples are shown in **Figure S3G-H**). Charge reversals at three conserved negatively charged residues in MAD1^41-62^ (E52K, E53K, E56K) abolished binding to Cyclin B1 (**Figure 2C-D**). To identify potential binding partners on Cyclin B1 for the MAD1 residues E52, E53, and E56, we mutagenized various clusters of positively charged residues on the surface of Cyclin B1, without however identifying a sufficiently penetrant mutant (**Table S1, Part III**). Collectively, these results indicate that MAD1 and Cyclin B1:CDK1 interact directly, and that the interaction is mediated primarily or exclusively by residues 41-62 of MAD1 and by Cyclin B1. In addition, a conserved acidic patch in this N-terminal region of MAD1 is essential for Cyclin B1 interaction (**Figure 2E**).

To assess the effect of inhibiting Cyclin B1-MAD1 interaction in cells, we generated doxycycline-inducible vsv-tagged MAD1-WT or MAD1-3EK HeLa cells and used these to create MAD1 knockouts via CRISPR/Cas9 (with a gRNA targeting Exon 3 to knockout MAD1α and MAD1β; **Figure 2F**). MAD1 localisation was then assessed in nocodazole-arrested cells, which demonstrated that MAD1-WT and MAD1-3EK displayed a similar localisation pattern in early prometaphase, but only the MAD1-WT was able to localise to the corona when it formed in late prometaphase (**Figures 2G**). At this stage, MAD1-3EK levels decreased at the kinetochore (**Figures 2G-H**), which is consistent with the fact that the BUB1-dependent pool of MAD1 is reduced by PP2A as mitosis progresses [31]. Therefore, a MAD1-3EK mutant, which is unable to bind directly to Cyclin B1, is also unable to localise to the corona in nocodazole-arrested cells. This confirms that Cyclin B1 is the scaffold that recruits MAD1 to this region of the kinetochore in human cells. When the corona pool is removed in MAD1-3EK cells, MAD1 kinetochore recruitment is reduced soon after nuclear envelope breakdown (mirroring the localisation and phosphorylation of its other kinetochore receptor, BUB1) [31, 32]. Note, that we also generated YFP-tagged MAD1 cells to visualise its localisation live. However, YFP-MAD1-WT and YFP-MAD1-3EK were both absent from the corona, which suggests that a large N-terminal tag affects MAD1 localisation to this region (**Figure S4**). This may be why removing the N-terminus of GFP or mCherry-MAD1 was not reported to affect MAD kinetochore localisation in previous studies [18, 20]. It is also important to note that the N-terminal vsv-tag on MAD1 is not detected at the corona by immunofluorescence (results not shown), suggesting that this region maybe buried in an interaction interface.

### Corona-localised MAD1 generates a robust SAC response

The ability of Cyclin B1 to recruit MAD1 to the corona could allow Cyclin B to generate the signal that inhibits its own degradation. However, it is unclear if corona-localised MAD1 can signal directly to the SAC and, if it can, how this differs from the conventional KNL1-BUB1-MAD1 pathway at the outer kinetochore. One major difference is that Cyclin B1-MAD1 localisation to the corona is insensitive to MPS1 inhibition after mitotic entry (**Figure 1E**), whereas MPS1 activity is continually required to phosphorylate KNL1 [33–36] and BUB1 [31, 37–41] to recruit MAD1 to the outer kinetochore. To investigate this major difference between the two pathways, we tested the response of MAD1-WT and MAD1-3EK cells to MPS1 inhibition. As expected [15], MAD1 was preserved on kinetochores following MPS1 inhibition with AZ-3146 after mitotic entry in MAD1-WT cells (**Figure 3A-C**). However, in stark contrast, a MAD1-3EK mutant that cannot bind the corona was completely lost from kinetochores under identical conditions (**Figure 3A-C**). This has considerable impact on the SAC, because MAD1-3EK cells are exquisitely sensitive to MPS1 inhibition in nocodazole, as demonstrated by the fact that these cells immediately exit mitosis at a dose of AZ-3146 that can be tolerated for several hours in MAD1-WT cells (**Figure 3D**). These data demonstrate that Cyclin B1-MAD1 becomes insensitive to MPS1 inhibition once the corona has been established, which subsequently allows the SAC to tolerate substantial reductions in MPS1 activity.

**Figure 3.**
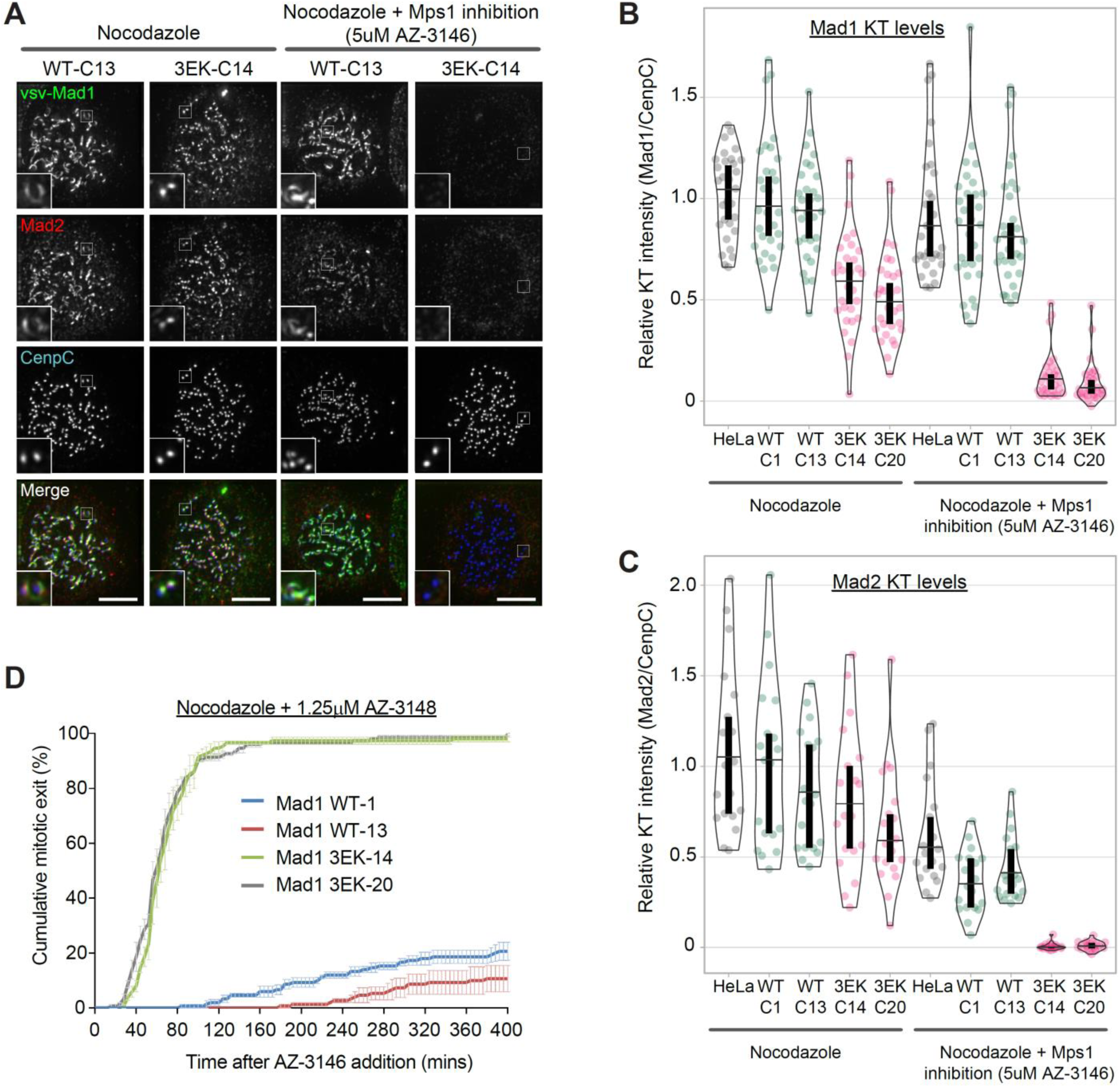
Inhibiting Cyclin B1-MAD1 interaction weakens the SAC. **A-C.** Immunofluorescence images (A) and quantifications (B,C) of MAD1 and MAD2 kinetochore levels in indicated cells lines arrested in nocodazole and treated with/without AZ-3146 for 30 min to inhibit MPS1. In all kinetochore intensity graphs, each dot represents a cell, horizontal lines indicate the median and vertical bars show the 95% confidence interval. 60 cells (MAD1) and 30 cells (MAD2) from 3 experiments. **D.** Duration of mitotic arrest in indicated cell lines arrested in nocodazole and then treated with 1.25 µM AZ-3146. Graph shows cumulative mean (±SEM) of 3 experiments, 50 cells per condition per experiment.

### Cyclin B scaffolds MAD1 at the corona to allow the SAC to tolerate MPS1 inhibition

As well as regulating MAD1 recruitment to the outer kinetochore, MPS1 activity is also needed to phosphorylate the C-terminal domain (CTD) of MAD1 and catalyse MCC assembly [37, 38, 42]. Therefore, we reasoned that MAD1 may still need to be phosphorylated by MPS1 to catalyse MCC assembly from the corona. Therefore, to probe this further, we raised a phospho-specific antibody to a key Thr716 MPS1-phosphorylation site on MAD1 and confirmed its specificity in cells (**Figure S5**). This antibody detects a strong signal at unattached kinetochores in RPE1 and HeLa cells, which is rapidly lost upon MPS1 inhibition (**Figures 4A and S5C**). Importantly, in nocodazole-arrested cells, although MAD1 decorates the whole corona, the MAD1-pT716 signal is restricted to the outer kinetochore around the KMN network (**Figure 4A and S5C**). This suggests that MPS1 has a limited zone of activity that is relatively confined to its anchor point on NDC80 [6, 7]. How then can the corona MAD1 support the SAC following MPS1 inhibition? We hypothesised that this MAD1, which is tethered to the corona via Cyclin B1 at its N-terminus, may be able to use its predicted coiled-coil to allow the CTD to reach the zone of MPS1 activity at NDC80 [14].

**Figure 4.**
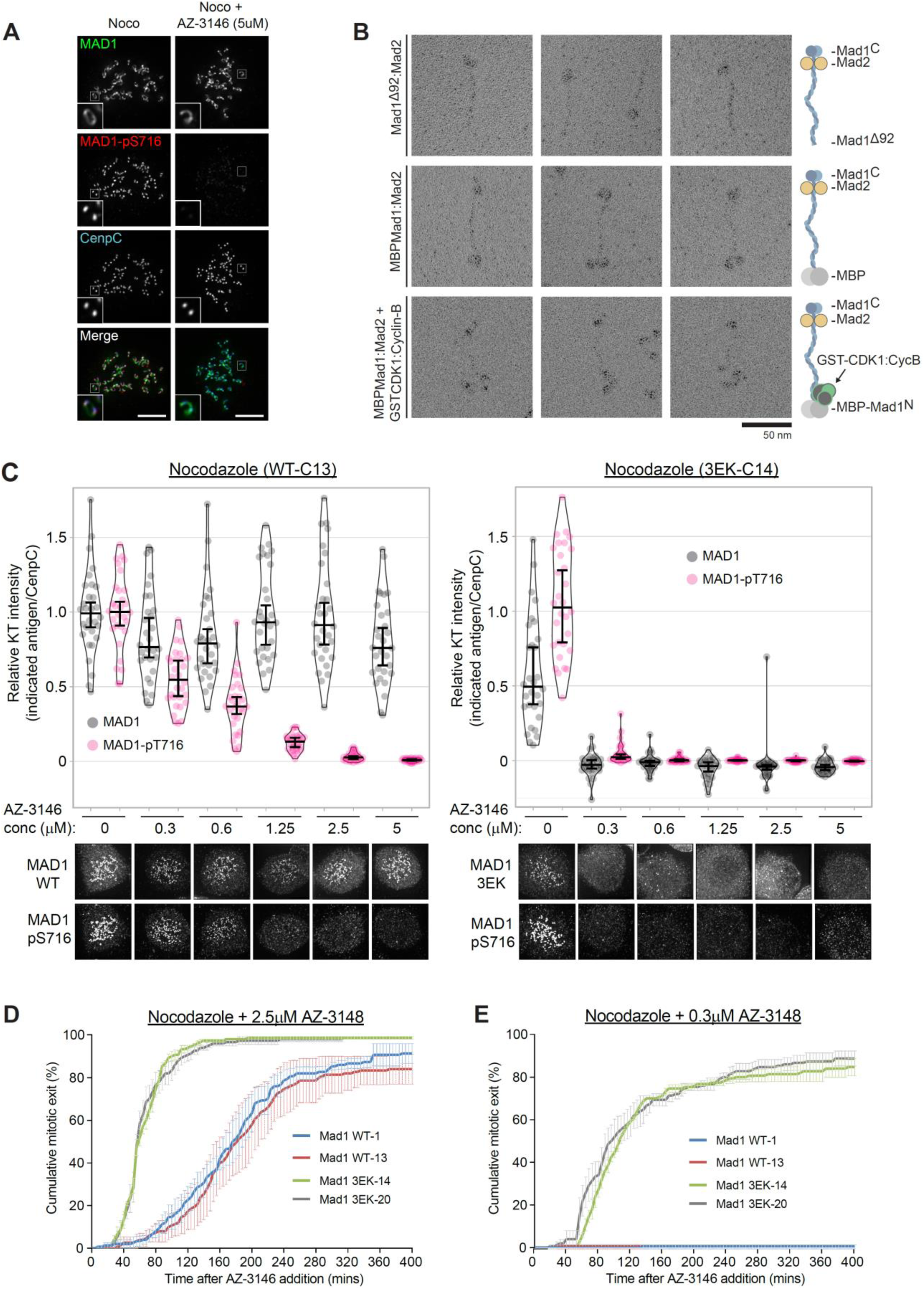
Cyclin B scaffolds MAD1 at the corona to allow the SAC to tolerate MPS1 inhibition. **A.** Immunofluorescence images of MAD1 and MAD1-pT716 kinetochore levels in nocodazole-arrested RPE1 cells. **B.** Electron micrographs of rotary-shadowed MAD193-718:MAD2 (top row), MBP-MAD1:MAD2 (middle row), and MBP-MAD1:MAD2 in complex with GST-CDK1:Cyclin B1 (bottom row). The scale bar indicates 50 nm. **C.** Quantifications (top) and corresponding immunofluorescence images (underneath) of kinetochore MAD1 and MAD1-pT716 levels in nocodazole-arrested of MAD1-WT-C13 and 3EK-C14 treated with different doses of AZ-3146 for 30 min. MG132 was included at the time of AZ-3146 addition to prevent mitotic exit. Each dot represents a cell, horizontal lines indicate the median and error bars show the 95% confidence interval. 30 cells from 3 experiments. **D**,**E.** Duration of mitotic arrest in indicated cell lines arrested in nocodazole and then treated with 2.5 μM AZ-3146 (D) or 0.3 μM AZ-3146 (E) or. Graph shows cumulative mean (±SEM) of 3 experiments, 50 cells per condition per experiment.

Therefore, we next set out to visualize the stable complex between MAD1:MAD2 and Cyclin B1:CDK1 by electron microscopy after low-angle metal shadowing. This highlighted the position of MAD2 near the C-terminal end of MAD1 and demonstrated that MAD1 adopted a thin elongated structure with an apparent length of ∼66 nm (**Figure 4B**). Addition of the globular MBP (43 kDa) allowed the N-terminal end of full-length MAD1 to be recognized within MAD1:MAD2 complexes (**Figure 4B**). We then inspected a SEC fraction containing Cyclin B1:CDK1 bound to MBP-MAD1:MAD2 and identified an additional density near the N-terminal MBP in a number of complexes (**Figure 4B**). Thus, CDK1:Cyclin B1 binds the very end of an elongated MAD1:MAD2 complex and the opposite end, which is a substrate for MPS1 [37, 38, 42], lies approximately 66 nm away. This observation suggests that corona-localised MAD1 can still be phosphorylated by MPS1 at the KMN network as long as the anchor point for Cyclin B1 is within ∼66 nm of NDC80.

We hypothesised that corona-localised MAD1 could therefore help the SAC to tolerate MPS1 inhibition because, despite the fact the KNL1-MELT and BUB1 are dephosphorylated, corona-MAD1 is still preserved at kinetochores to respond to low levels of MPS1 activity. To test this hypothesis, we stained for MAD1 and MAD1-pT716 in nocodazole-arrested MAD1-WT or MAD1-3EK cells treated with a range of doses of the MPS1 inhibitor AZ-3146. **Figures 4C and S6A** show that total MAD1 protein and MAD1-pT716 are removed together from kinetochores at very low doses of MPS1 inhibitor in MAD1-3EK cells. However, the MAD1 is preserved at kinetochore following MPS1 inhibition in WT cells, which allows MAD1 phosphorylation to persist until much higher doses of AZ-3146. Quantifying mitotic exit over this dose range shows that the duration of mitotic arrest correlates with the amount of MAD1-pT716 at kinetochores (**Figure S6**). A persistent mitotic arrest is seen whenever MAD1-pT716 is detected on kinetochores, which equates to ≤ 0.3 µM AZ-3146 in MAD1-3EK cells and ≤ 2.5 µM AZ-3146 in MAD1-WT cells. In fact, the length of mitotic arrest is very similar in MAD1-3EK and MAD1-WT cells treated with 0.3 µM and 2.5 µM AZ-3146, respectively (**Figure 4D-E**), and these two conditions give almost identical levels of MAD1-pT716 (**Figures 4C and S6A**). The actual MAD1-pT716 signal in these two situations is just detectable on kinetochores, which presumably reflects a minimal sufficient amount to generate a SAC response.

## DISCUSSION

We show here that Cyclin B1 allows the SAC to tolerate substantial MPS1 inhibition by tethering MAD1 at the corona in a manner that is MPS1-independent once the corona has been established. Although this distinguishes corona-localised MAD1 from the canonical pool at the KMN network, it most likely behaves similarly in all other respects, because MPS1 phosphorylation at the C-terminus of MAD1 is most likely still required to catalyse MCC assembly [37, 38, 42]. We speculate that the thin elongated structure of MAD1 facilitates this process by providing the necessary reach to orient MAD2 and the MAD1 CTD towards MPS1 at the KMN network.

This link between corona MAD1 and the kinetochore SAC signal explains why end-on microtubule attachments rapidly silence the SAC in human cells: microtubules compete with MPS1 for NDC80 binding [6, 7], and in addition, they allow dynein motors to remove corona-localised MAD1 away from the zone of MPS1 activity [8–11]. Both components are necessary to fully extinguish the SAC because preventing either MPS1 or MAD1 removal from kinetochores is sufficient to also prevent SAC silencing [12, 13]. The final model is presented in figure 5.

**Figure 5.**
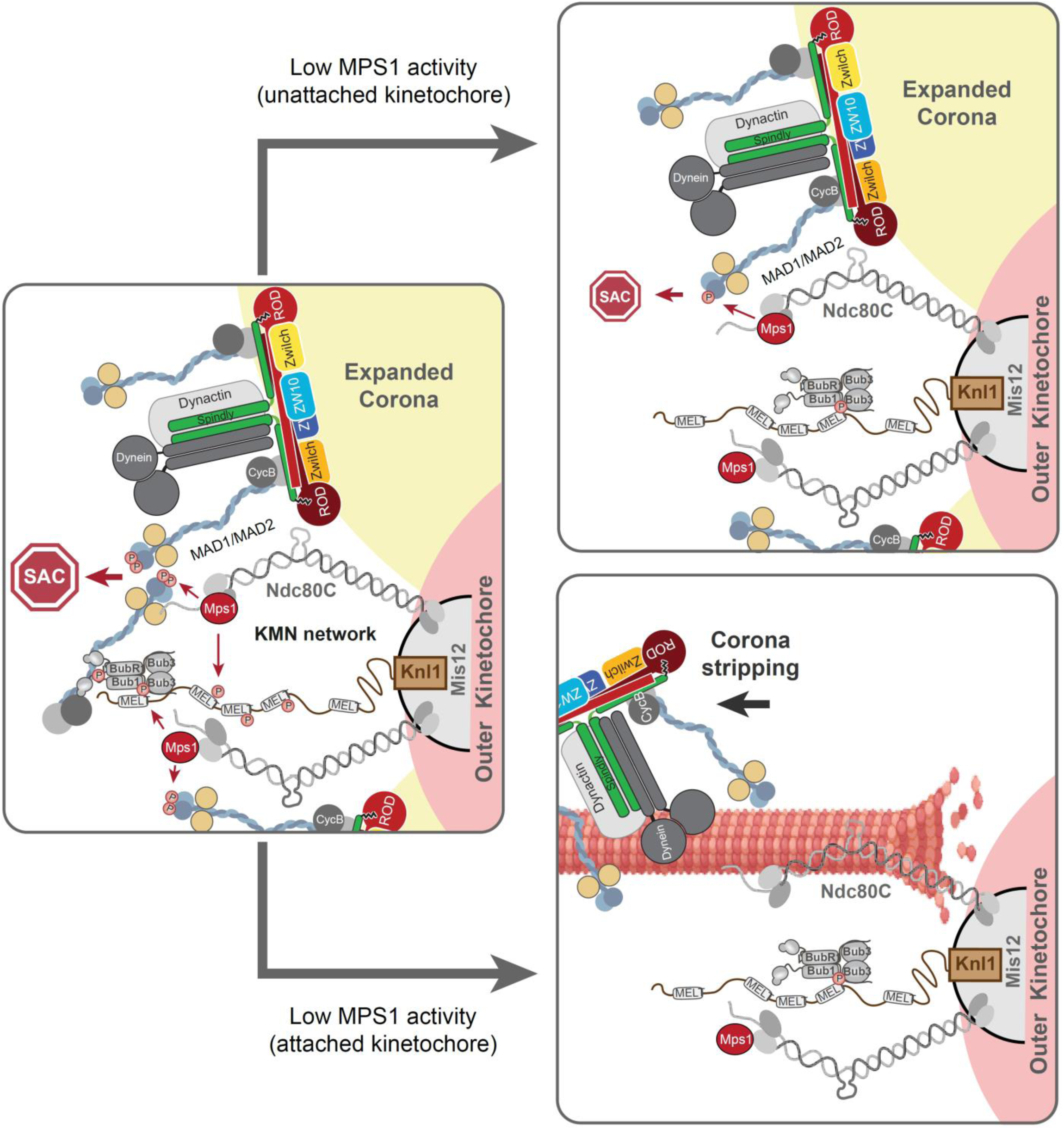
Model for how Cyclin B1-MAD1 at the corona helps the SAC to respond to low levels of MPS1 activity. **A.** On unattached kinetochore MAD1 is recruited via MPS1-phosphorylated BUB1 at the KMN network and Cyclin B1 at the corona. **B.** If MPS1 activity is lowered without microtubule attachment, then the BUB1-dependent pool of MAD1 is removed at the KMN network but the corona MAD1 pool persists to respond to low level of MPS1 activity and generate a SAC signal. **C.** When microtubules attach then MPS1 activity is lowered and the corona is stripped to remove all MAD1 and silence the localised SAC signal.

There are two other recent studies that also demonstrate that MAD1-Cyclin B1 interaction is important during mitosis, but for different reasons. Alfonso-Peres et al demonstrated that knockdown of MAD1, or removal of its first 100 amino acids, causes a partial SAC defect in nocodazole and reduces the amount of Cyclin B1 and MPS1 on kinetochores (by approximately 75% and 50%, respectively) [18]. It should be noted, however, that the concentration of nocodazole used in these studies (0.3μM) may be insufficient to fully depolymerise microtubules [43], which complicates interpretations about direct effects on the SAC and on MAD1/Cyclin B1/MPS1 localisation to unattached kinetochores. Furthermore, removal of the N-terminal region of MAD1 has previously been shown to affect MCC assembly from the nuclear pore [20], which may also have contributed to the observe SAC defects. Nevertheless, the authors put forward an important hypothesis by proposing that kinetochore Cyclin B1/CDK1 may drive localised CDK1 activity to support the SAC; for example, by increasing CDK1-mediated phosphorylation of MPS1 on Ser281 to enhance MPS1 localisation (see accompanying paper by the same groups [44]). Although kinetochore localised Cyclin B1 has not so far been demonstrated to drive CDK1 substrate phosphorylation at kinetochores, it will be important to test this hypothesis in future because, if it does, then this has the potential to impact on many processes, including the SAC. As well as MPS1 localisation, CDK1 positively and negatively regulates a number of other key enzymes at the kinetochore (for review see: [45]), and therefore these substrates may change dramatically upon kinetochore-microtubule attachment when Cyclin B1/MAD1 is stripped away down microtubules. Furthermore, the removal of the corona itself may depend on localised CDK1 inactivation because acute inhibition of CDK1 is known to cause premature corona detachment [27, 46]. To address the potential importance of such localised kinetochore CDK1 regulation, however, we believe it will be crucial to first identify the other receptor(s) for Cyclin B1 at the corona so that this pool can then be fully removed from unattached kinetochores.

Very recent data from Jackman et al has also demonstrated the importance of Cyclin B1-MAD1 interaction during mitosis, but this time in the release of MAD1 from the nuclear pore [19]. In this study, preventing the interaction delays MAD1 accumulation at kinetochores until late prophase, as well as weakening the SAC and enhancing the level of chromosomal instability (CIN). The authors use an elegant approach to mutate two key acidic residues in MAD1 (E53K/E56K) at the endogenous locus of RPE cells by CRISPR/Cas9. Although this reduces the amount of Cyclin B1 that co-precipitates with MAD1, it is possible that the additional E52K mutation included in our 3EK mutant may produce a more penetrant phenotype, since all three glutamates lie on the same face of the predicted helix in MAD1 [19]. Nevertheless, the subsequent results on MAD1 dissociation from the nuclear pore are entirely consistent with our data, since these focus on an earlier stage of mitosis and the authors do not examine the effects later in prometaphase at unattached kinetochores. It is likely, therefore, that Cyclin B1-MAD1 functions at the nuclear pore in early mitosis and then again at the corona following nuclear envelope breakdown to safeguard chromosome segregation and prevent CIN. It will be interesting to determine whether this interaction is commonly perturbed in cancer cells that frequently missegregate their chromosomes to become chromosomally unstable.

In summary, this study reveals how the main mitotic kinase, Cyclin B1/CDK1, plays a key role in scaffolding the SAC machinery to the corona. Considering that Cyclin B1 is ultimately degraded by the APC/C once the SAC has been silenced, this important scaffolding function most likely helps to ensure the SAC cannot be re-established following anaphase onset.

## ACKNOWLEDGEMENTS

This study was funded by Cancer Research UK (C47320 to A.T.S. that also funds L.A), a Leng Charitable Trust PhD studentship to M.R., and by the Max Planck Society and the European Research Council (ERC) Advanced Investigator Grant RECEPIANCE (proposal 669686) to A.M. Microscopy was carried out at the Dundee Imagine Facility. We are grateful to Iain Cheeseman for providing the inducible MAD1α and β knockout line, Stephen Taylor for the HeLa Flp-in line, and Jane Endicott and Martin Noble for recombinant Cyclin B1 and CDK1 constructs.

## AUTHOR CONTRIBUTIONS

A.T.S discovered the MAD1-Cyclin B1 connection whilst working in the lab of G.K. (figures 1A-C, 1F-G). L.A. performed most other cell biology experiments, with important contributions from M.R. The *in vitro* biochemistry was performed by Y.L. and the EM of MAD1 by P.H.V., under the supervision of A.M. A.T.S. wrote the manuscript with comments from all authors.

## MATERIALS AND METHODS

### Cell culture and reagents

RPE1 were purchased from ATCC and HeLa Flp-in cells were a gift from S Taylor (University of Manchester, UK) [47]. The RPE1 Cyclin B1-EYFP cells have been published previously [48], as have the U2OS with LacO array on chromosome 1 [49]. All cells were authenticated by STR profiling (Eurofins) and screened every 4-8 weeks to ensure they were mycoplasma free. Cells were cultured in DMEM supplemented with 9% FBS and 50 µg/ml penicillin/streptomycin. Except during fluorescence time-lapse analysis, when they were cultured in Leibovitz’s L-15 media (900 mg/L D+ Galactose, 5mM Sodium Pyruvate, no phenol red). Doxycycline (1µg/ml), STLC (S-Trityl-L-cystiene: 10 μM) and thymidine (2 mM) were purchased from Sigma-Aldrich, nocodazole (3.3 µM) from Millipore, puromycin and hygromycin B from Santa Cruz Biotechnology, MG132 (10 µM) from Selleckbio, AZ-3146 (at indicated concentrations) from Axon, Rapamycin (100nM) from LC Laboratories, and reversine (at indicated concentrations) from Cayman Chemicals.

### Generation of knockout cells lines by CRISPR/Cas9 gene editing

To generate MAD1α knockout cells (i.e. MAD1β cells), a guide RNA targeting exon 4 of Mad1 (CCGCTCCACCTGGATGAGGTGGG) was cloned into a lentiviral vector that co-expresses Cas9 and a puromycin resistance marker (pLentiCRISPRv2; addgene #52961) to create pLentiCas9-g4-MAD1. Cyclin B-YFP-FKBP HeLa Flp-in cells (generated by CRISPR/Cas9-mediated homologous recombination) were transfected with pLentiCas9-g4 plasmid and selected in puromycin to obtain single cell clones. These were screened for the absence of nuclear MAD1 by immunofluorescence, since only MAD1α, and not MAD1β, is localised to the nucleus in interphase [21], and 2 clones were subsequently validated (C13 and C24: **Figure S2**). Note, the original aim was to target endogenous MAD1 and replace exon 4 with an FRB cassette to lose Cyclin B interaction and regain it upon rapamycin addition (to induce Cyclin B-YFP-FKBP interaction). The knock-in gene-editing was however unsuccessful, therefore MAD1α knockouts were used instead. To generate doxycycline-inducible vsv or YFP-MAD1-WT and -3EK cell lines HeLa Flp-in cells were transfected with Mad1 in pCDNA5/FRT/TO vector (Invitrogen) together with the FLP recombinase, pOG44 (Invitrogen) using Fugene HD (Promega) according to the manufacturer’s protocol. Stable transfectants were selected in media containing 200 μg/mL hygromycin-B. Subsequently, to knock-out endogenous Mad1, these cells were transfected with a guide RNA targeting exon 3 of Mad1 (CTTCATCTCTCAGCGTGTGGAGG) in pLentiCRISPRv2 for 24h and thereafter selected in puromycin for a further 24h. Cells were then cultured continually in the presence of Dox to maintain viability after knock-out of endogenous Mad1 by inducing expression of vsv- or YFP-Mad1 WT or 3EK. Individual clones were isolated and screened for loss of endogenous Mad1 by western analysis for YFP-tagged Mad1 (**Figure S4)** and 10 days after removal of Dox for vsv-tagged Mad1 (**Figure 2F)**. 2 clones for each construct were validated and used subsequently (vsv-Mad1 WT: C1 & 13; vsv-Mad1 3EK: C14 & C20; YFP-Mad1 WT: C5 & C19; YFP-Mad1 3EK: C6 & C18).

### Antibodies

The following primary antibodies were used for immunofluorescence imaging (at the indicated final concentration diluted in 3% BSA in PBS): chicken α-GFP (ab13970 from Abcam, 1:5000), guinea pig α-Cenp-C (BT20278 from Caltag + Medsystems), human ACA (90C-CS1058 from Fitzgerald, 1:2000), rabbit Cyclin B1 (#12231S from Cell signalling technology, 1:1000), mouse Mad1 (clone BB3-8, MABE867 from Millipore, 1:1000), mouse Tubulin (clone B-5-1-2 from Sigma, 1:5000), rabbit ZW10 (ab21582 from abcam, 1:1000), MAD2 (A300-301A-T from Bethyl, 1:1000). The MAD1-pT716 used in this study (MAD1-pT716-p1) was a custom rabbit polyclonal phospho-specific antibody generated by Biomatik. Secondary antibodies used for immunofluorescence were highly-cross absorbed goat, α-chicken, α-rabbit, α-mouse or a-guinea pig coupled to Alexa Fluor 488, Alexa Fluor 568, or Alexa Fluor 647 (Thermo Fischer). The primary antibodies used for Western blotting were Actin (A2066 from Sigma, 1:5000), FKBP12 (Clone H-5 from Santa Cruz, 1:250), mouse Mad1 (clone BB3-8, MABE867 from Millipore, 1:5000), tubulin (clone 5-B-1-2 from Sigma, 1:5000), GST (clone 8-326, MA4-004 from Thermo Fischer, 1:1000) and Cyclin B (C8831 from Sigma, 1:1000). The secondary antibodies used for Western blotting were goat ⍰ -mouse IgG HRP conjugate (170-6516 from Bio-Rad; 1:2000) and goat ⍰ -rabbit IgG HRP conjugate (170-6515 from Bio-Rad; 1:5000).

### Time-lapse analyses

For fluorescence imaging, cells were imaged in 8-well chamber slides (ibidi), on either a Zeiss Axio Observer 7 with a CMOS Orca flash 4.0 camera or a Deltavision Elite equipped with Photometrics CascadeII:1024 EMCCD or CoolSNAP HQ (Photometrics) camera. The objectives used for fluorescent imaging where either 20x/0.8 NA or 40x/1.3NA. For brightfield imaging, cells were imaged in a 24-well plate in DMEM on a Zeiss Axiovert 200M using Hamamatsu ORCA-ER camera and controlled by Micro-manager software (open source: https://miro-manager.org/), or a Zeiss Axio Observer 7 as detailed above. The air objectives used for brightfield imaging were either 10x/0.5 NA or a 20x/0.4 NA. Mitotic exit was defined by cells flattening down in the presence of nocodazole and Mps1 inhibitor.

### Immunofluorescence

Cells, plated on High Precision 1.5H 12-mm coverslips (Marienfeld), were treated and then pre-extracted with 0.1% Triton X-100 in PEM (100 mM Pipes, pH 6.8, 1 mM MgCl_2_ and 5 mM EGTA) for 1 minute before addition of 4% paraformaldehyde (PFA) in PBS for 10 min. Experiments involving MAD1-pT716 were not pre-extracted and fixed directly in 4% PFA Coverslips were washed with PBS and blocked with 3% BSA in PBS + 0.5% Triton X-100 for at least 30 min. Thereafter coverslips were incubated with primary antibodies overnight at 4°C, washed with PBS and incubated with secondary antibodies plus DAPI (4,6-diamidino2-phenylindole, Thermo Fischer) for an additional 2-4 hours at room temperature in the dark. Coverslips were then washed with PBS and mounted on glass slides using ProLong antifade reagent (Molecular Probes). All images were acquired on a DeltaVision Core or Elite system equipped with a heated 37°C chamber, with a 100x/1.40 NA U Plan S Apochromat objective using softWoRx software (Applied precision). Images were acquired at 1x1 binning using a CoolSNAP HQ or HQ2 camera (Photometrics) and processed using softWorx software and ImageJ (National Institutes of Health). All immunofluorescence images displayed are maximum intensity projections of deconvolved stacks and were chosen to most closely represent the mean quantified data.

### Image quantification

For quantification of immunostainings, all images to be compared directly were acquired with identical illumination settings. An ImageJ macro was used to threshold and select all kinetochores and all chromosome areas (excluding kinetochores) using the DAPI and anti-kinetochore antibody channels, as previously [50]^37^. This was used to calculate the relative mean kinetochore intensity of various proteins ([kinetochores-chromosome arm intensity (test protein)] / [kinetochores-chromosome arm intensity (CENP-C)]. For the quantification of kinetochore localisation on attached vs unattached kinetochore, cells were arrested in STLC and MAD1 intensity used to define attached (MAD1 negative) or unattached (MAD1 positive) kinetochores. The signal in the MAD1 and Cyclin B channel was then expressed as a percentage of CenpC on these individual kinetochores (after normalising for surrounding background intensities). Graphs were generated using either Graphpad or PlotsOfData [51].

### DNA cloning, Protein Expression and Purification

Constructs for the recombinant expression of GST-CDK1:Cyclin B1 and CDK1-Cyclin B1^165-413^ complexes were gifts of Jane Endicott and Martin Noble, and the complex was expressed and purified as previously described [52]. The MBP-MAD1:MAD2 complex and MAD1^93-718^:MAD2 complex were expressed and purified as described [37]. DNA sequences encoding MAD1 N-terminal fragments were amplified from a full-length MAD1 synthetic gene encoding the HsMAD1 sequence [37] by the polymerase chain reaction (PCR) and sub-cloned into the previously generated pETDuet-MBP8His vector [53]. All the mutant versions of recombinant proteins were produced by QuickChange Mutagenesis kit to mutate the plasmid DNA (Agilent Technologies). The MAD1 fragments and Cyclin B1 were transformed into BL21(DE3) Rosetta 2 competent cells. Cells were grown in Terrific broth at 37°C to an OD_600_ of about 1.2. Protein expression was induced by addition of 1 mM IPTG at 37°C, and cells were further allowed to grow for 5 hours. Cell pellets were resuspended in binding buffer (20 mM Tris/HCl pH 8.0, 500 mM NaCl, 5% (v/v) glycerol, 1 mM EDTA, 1 mM TCEP), lysed by sonication and cleared by centrifugation at 10000x*g* for 45 min. The cleared supernatant was purified through Histrap HP and gel-filtered on a Superdex 200 10/300 columns (GE Healthcare). The LacI-GFP-MAD1 fragments were generated by PCR amplification of MAD1 and insertion into the LacI-GFP vector [54]. To create the guide RNA-resistant MAD1-WT and 3EK plasmids, gene blocks were synthesised that encode for amino acids 1-97 of MAD1, with or without mutations to change E52K, E53K and E56K (denoted as 3EK), and containing additional silent mutations in the gRNA sequences. These gene blocks were inserted into full length MAD1 by Gibson assembly to replace the region encoding for amino acids 1-97.

### Analytical Size-Exclusion Chromatography

All samples were eluted under isocratic conditions at 4° C in SEC buffer (20 mM Tris, 150 mM NaCl, 1 mM TCEP) at a flow rate of 0.1 ml/min. The elution profiles of proteins were monitored at 280 nm. To form the complex, proteins were mixed at 1:1 molar ration and a typical concentration of 10 µM) and incubated for 1 hour on ice. The loading volume for each injection was 25 µl and the final concentration of each protein is 5 µm. SDS-PAGE, followed by Coomassie staining, was used to detect proteins.

### Low-angle metal shadowing and electron microscopy

Fractions from size-exclusion chromatography containing protein complexes of interest were diluted 1:1 with spraying buffer (200 mM ammonium acetate and 60% glycerol) and air-sprayed onto freshly cleaved mica pieces (V1 quality, Plano GmbH). Specimens were mounted and dried in a MED020 high-vacuum metal coater (Bal-tec). A Platinum layer of approximately 1 nm and a 7 nm Carbon support layer were evaporated subsequently onto the rotating specimen at angles of 6-7° and 45° respectively. Pt/C replicas were released from the mica on water, captured by freshly glow-discharged 400-mesh Pd/Cu grids (Plano GmbH), and visualized using a LaB_6_ equipped JEM-1400 transmission electron microscope (JEOL) operated at 120 kV. Images were recorded at a nominal magnification of 60,000x on a 4k x 4k CCD camera F416 (TVIPS), resulting in 0.18 nm per pixel. Representative particles were manually selected using EMAN2 [55].

---

**Table S1 MAD1 and Cyclin B constructs used for *in vitro* studies**

**Part I.**
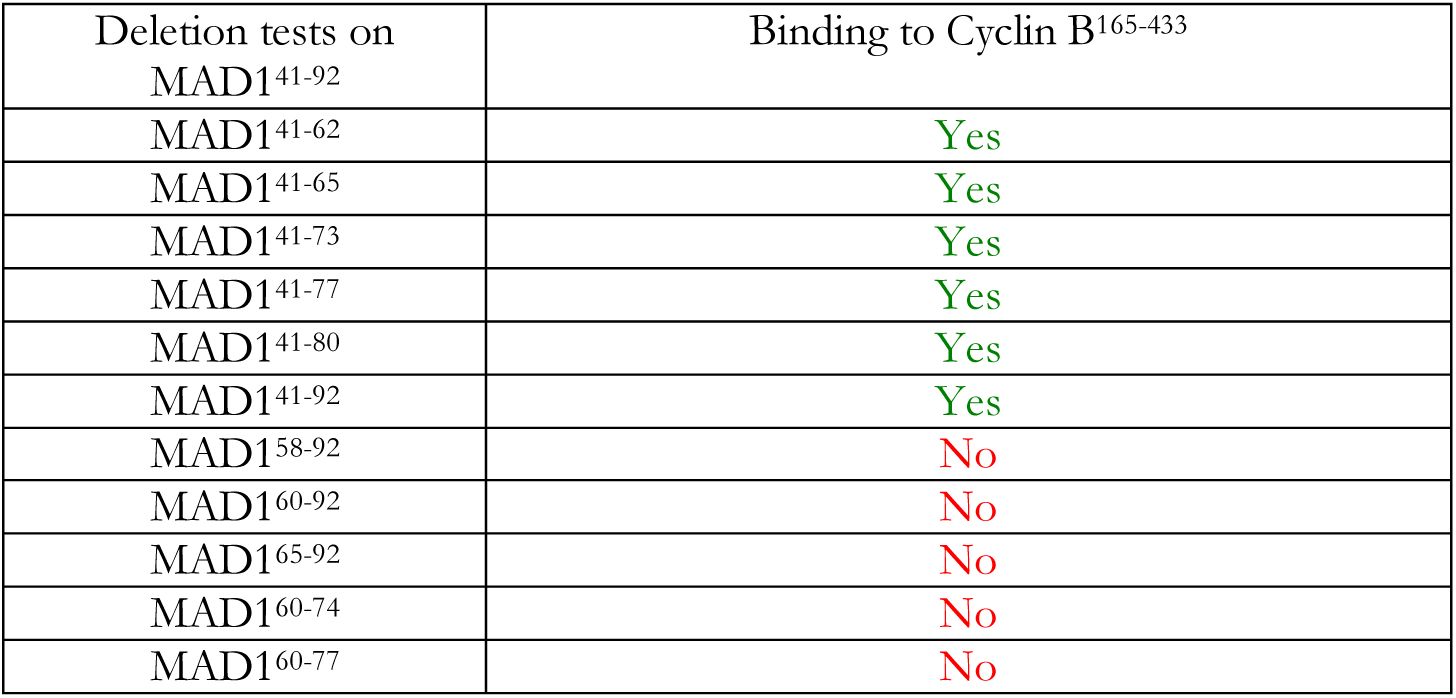
Identification of a minimal Cyclin B1-binding region in MAD1.

**Part II.** Mutagenesis of MAD1^41-62^ to identify Cyclin B1-binding determinants.

| MAD1<sup>41-62</sup><br>Mutation Group | Mutations | Binding to Cyclin B<sup>165-433</sup> |
| --- | --- | --- |
| M1 | Q42A, Q44A, Q46A,<br>Q47A, Q50A, Q57A | Yes |
| M2 | Q57A, S60A, S62A | Yes |
| M3 | E52K, E53K, E56K | No |
| M4 | Y45A, M49A, L51A,<br>I58A | Yes |

**Part III.** Mutagenesis of Cyclin B1 to identify MAD1-binding determinants.

| Cyclin B1<sup>165-433</sup><br>Mutation Group | Mutations | Binding to MBP-Mad1<sup>41-62</sup> |
| --- | --- | --- |
| M1 | K171E | Yes |
| M2 | K214E, R216E | Yes |
| M3 | R306E, R307E, K310E | Yes |
| M4 | K395E, K407E, K410E | Yes |
| M1 + M2 + M4 | K171E, K214E, R216E,<br>K395E, K407E, K410E | Yes |
| M5 | R187E, K189E, R194E | Yes |
| M6 | K240E, K241E | Yes |
| M7 | K287E, R290E | Yes |
| M6 + M7 | K240E, K241E, K287E,<br>R290E | Yes |
| M5 + M7 | R187E, K189E, R194E,<br>K287E, R290E | Yes |

**Figure S1.**
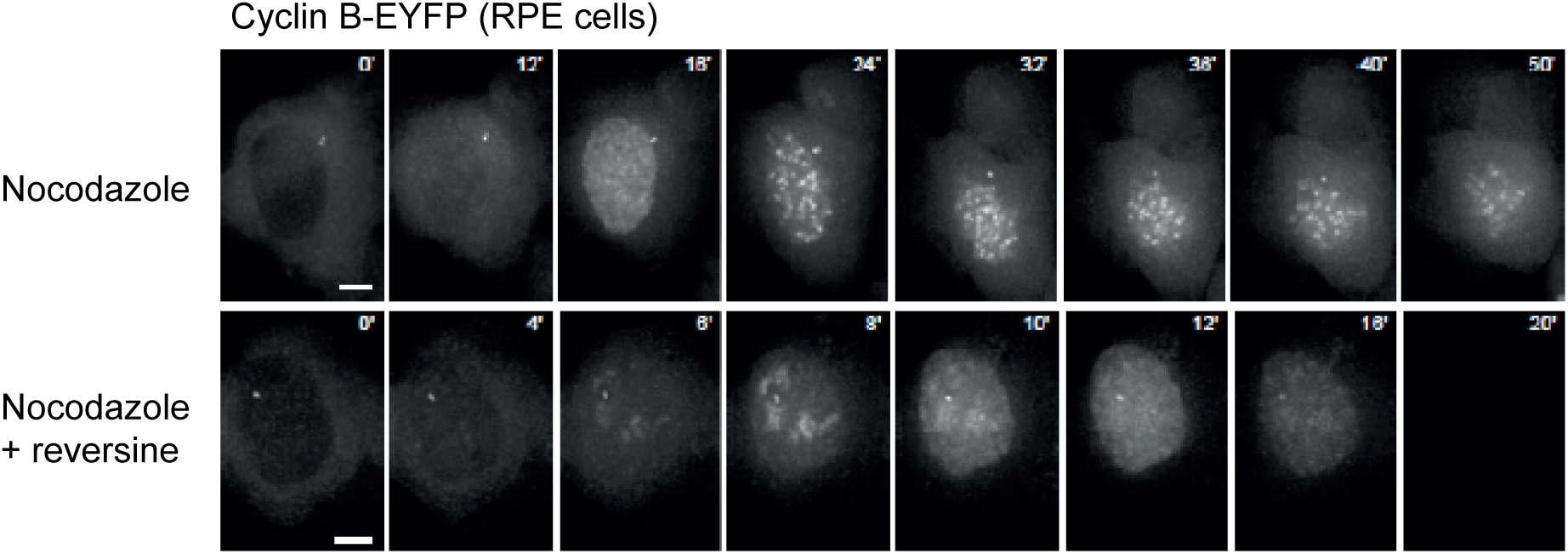
MPS1 is needed to establish Cyclin B at unattached kinetochores. Stills from movie of Cyclin B1-EYFP RPE1 cells enter mitosis and in the presence of nocodazole with or without 500nM Reversine.

**Figure S2.**
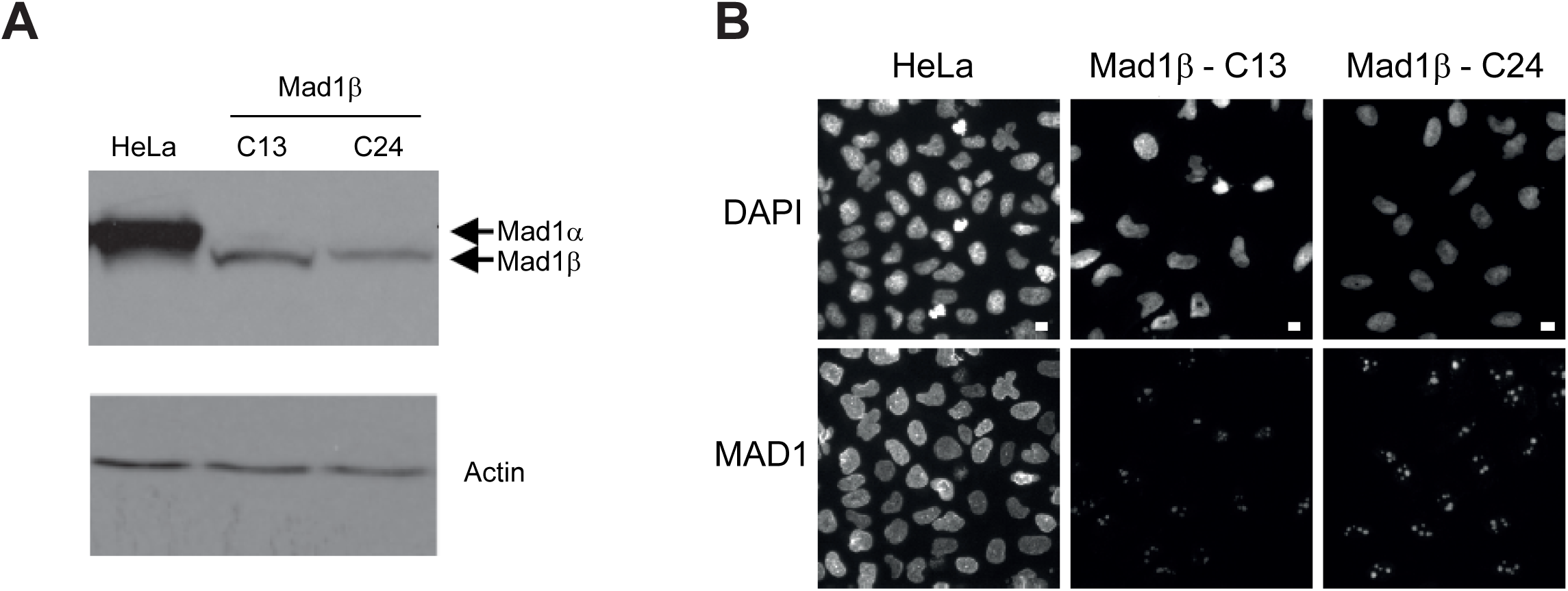
Validation of Mad1α knockout cell clones. **A**,**B.** Western blot (A) and immunofluorescence images (B) of MAD1 in control on MAD1α knockout HeLa cells (2 clones: MAD1α-C13 and C24).

**Figure S3.**
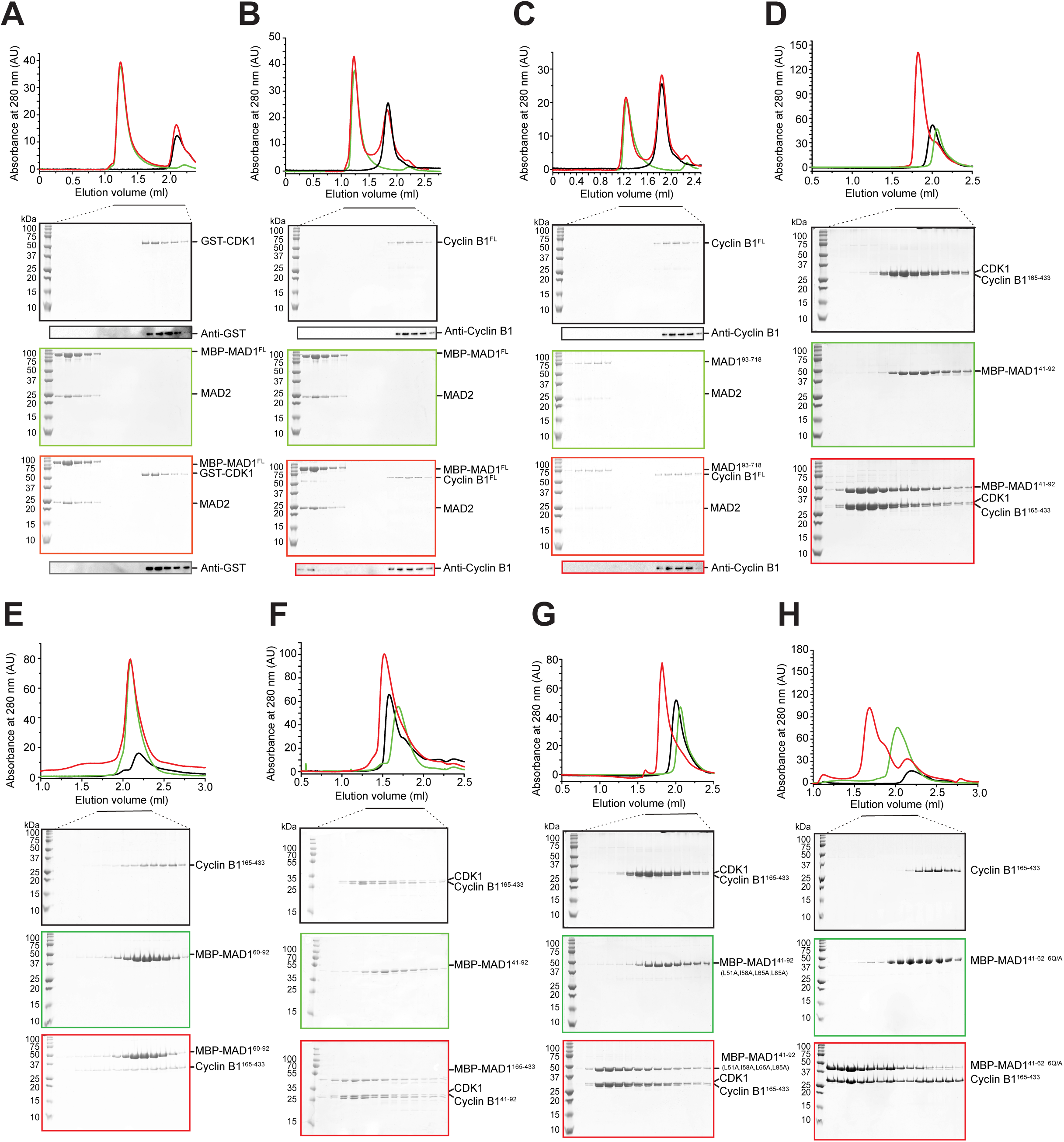
Cyclin B1, not CDK1, interacts with the MAD1:MAD2 complex via an acidic patch in the MAD1 N-terminus. **A-C.** Elution profiles and SDS-PAGE for SEC runs on a Superose 6 Increase 5/150 GL column of (A) GST-CDK1 complex (black), MBP-MAD1:MAD2 (green), and their combination (red); (B) Cyclin B1 (black), MBP-MAD1:MAD2 (green), and their combination (red); (C) Cyclin B1 (black), MBP-MAD193-718:MAD2 (green), and their combination (red). Western blots were obtained with duplicates of the shown SDS-PAGE gels, and obtained with the same elution fractions. In **A** to **C**, Western blots were performed from separate SDS-PAGE with the same elution fractions. In **B** and **C**, the same Cyclin B1FL elution profiles, SDS-PAGE, and western blots are shown as controls. **D.** Elution profiles and SDS-PAGE for SEC runs on a Superose 6 Increase 5/150 GL column of CDK1:Cyclin B1165-433 complex (black), MBP-MAD141-62 (green), and their combination (red). **E.** Elution profiles and SDS-PAGE for SEC runs on a Superose 6 Increase 5/150 GL column of Cyclin B1165-433 complex (black), MBP-MAD160-92 (green), and their combination (red). The same Cyclin B1165-433 elution profile and SDS-PAGE as in Figure 2C are shown as controls. **F.** Elution profiles and SDS-PAGE for SEC runs on a Superdex 200 Increase 3.2/300 column of GST-CDK1:Cyclin B1165-433 complex (black), MBP-MAD141-62 (green), and their combination (red). The same CDK1:Cyclin B1165-433 elution profile and SDS-PAGE as in Figure 2B are shown as controls. **G.** Elution profiles and SDS-PAGE for SEC runs on a Superdex 200 Increase 5/150 GL column of CDK1:Cyclin B1165-433 complex (black), MBP-MAD141-92 carrying L51A, I58A, L65A, L85A mutations (green), and their combination (red). **H.** Elution profiles and SDS-PAGE for SEC runs on a Superdex 200 Increase 5/150 GL column of CDK1:Cyclin B1165-433 complex (black), MBP-MAD141-6216carrying the Q42A, Q44A, Q46A, Q47A, Q50A, Q57A mutations (abbreviated as 6QA, green), and their combination (red). The same Cyclin B1165-433 elution profile and SDS-PAGE as in Figure 2D are shown as controls

**Figure S4.**
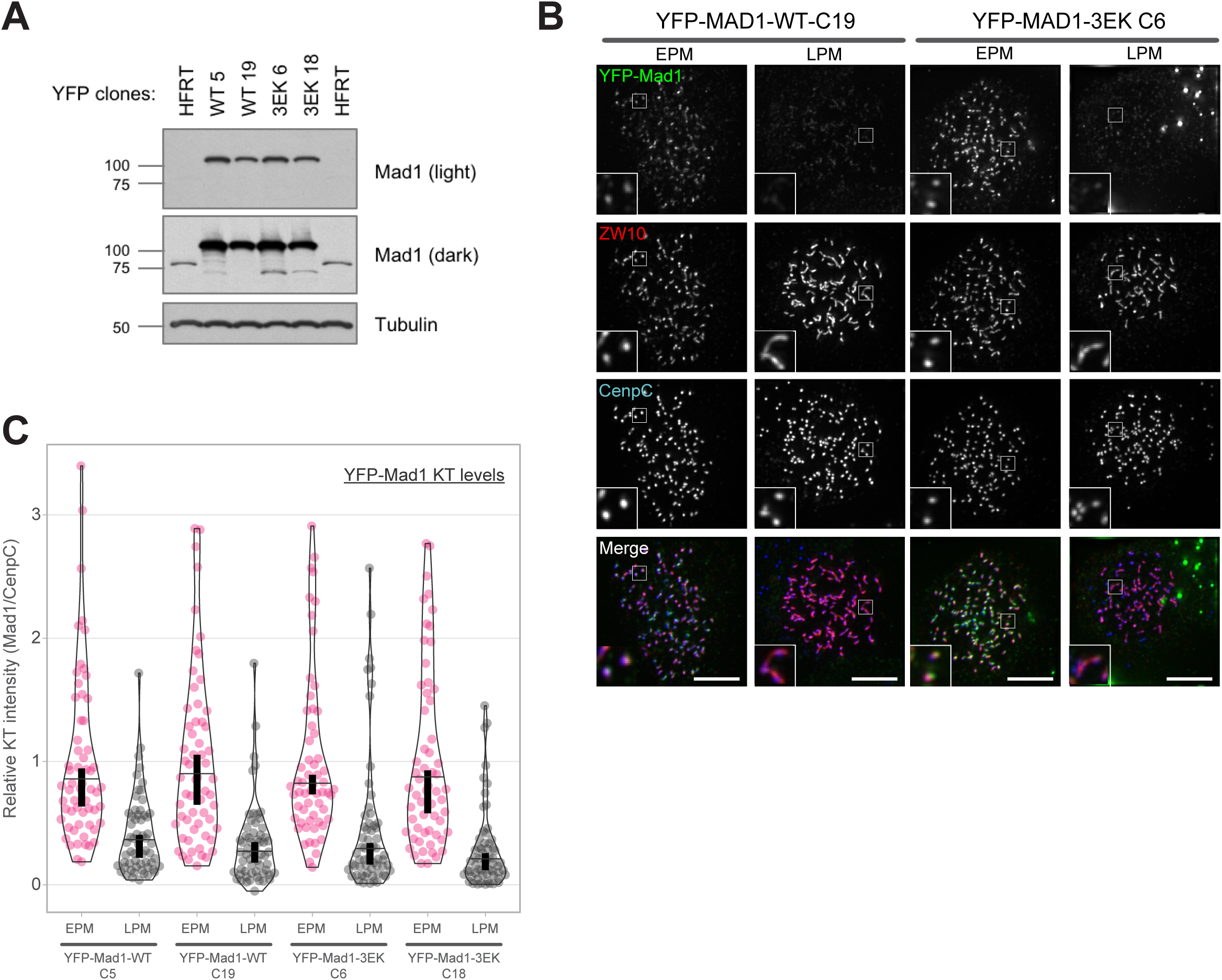
YFP-MAD1-WT and YFP-MAD1-3EK do not localise to the corona. **A.** Western blot analysis of indicated YFP-MAD1-WT and YFP-MAD1-3EK clones treated with doxycycline. **B.** Immunofluorescence images showing MAD1 and ZW10 kinetochore levels in nocodazole-arrested YFP-MAD1-WT-C19 AND 3EK-C6 cells just after nuclear envelope breakdown (early prometaphase - EPM) or later in mitosis when the chromatin is condensed (late prometaphase - LPM). **C.** Quantification of MAD1 kinetochore localisation from cells treated as in (B). In all kinetochore intensity graphs, each dot represents a cell, horizontal lines indicate the median and vertical bars show the 95% confidence interval. 60 cells from 3 experiments.

**Figure S5.**
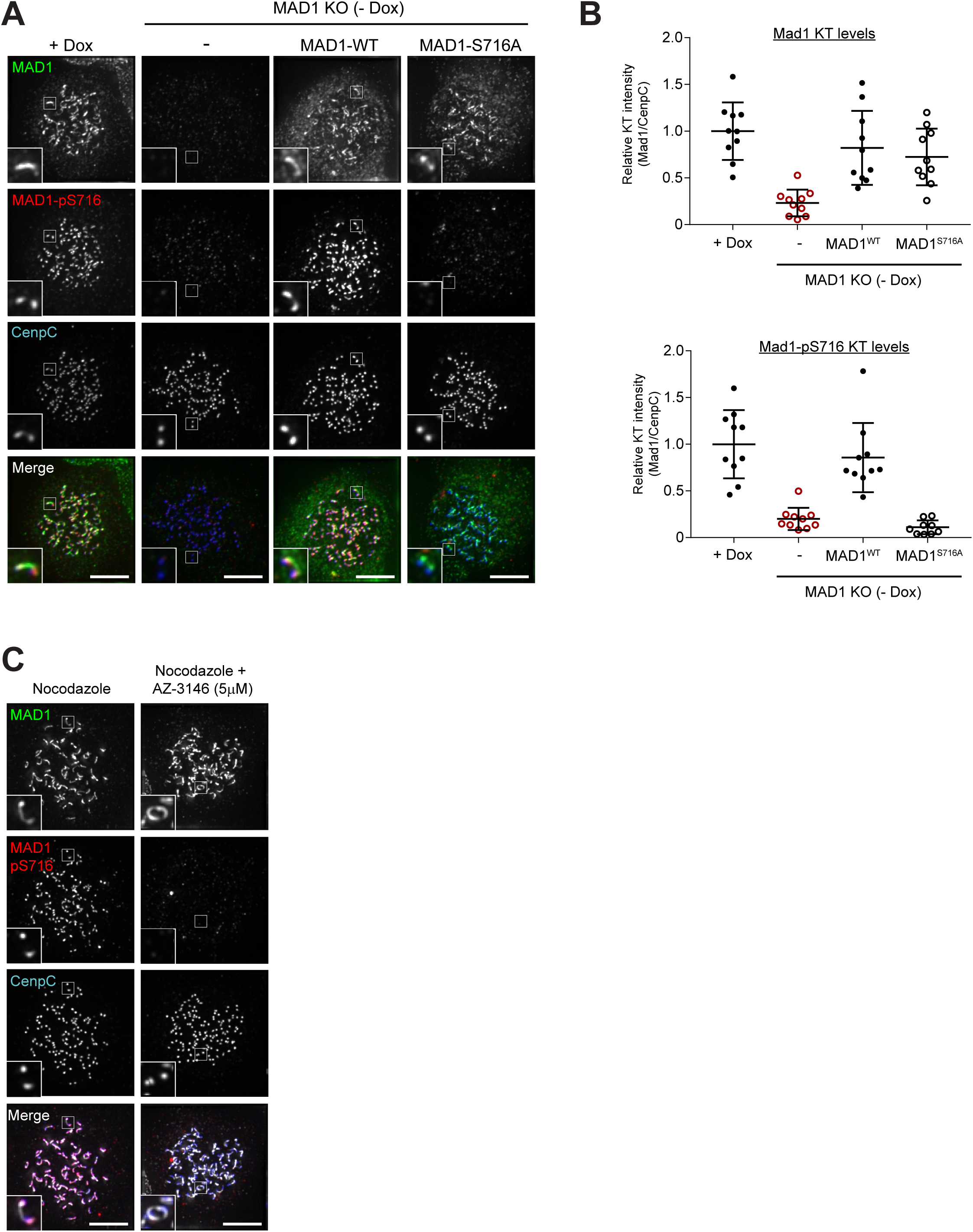
Validating the MAD1-pT716 antibody in cells. **A**,**B.** Immunofluorescence images (A) and quantification (B) of MAD1 and MAD1-pT716 kinetochore levels in dox-inducible vsv-MAD1-WT cells with doxycycline removed (-Dox) and MAD1-WT or MAD1-T716A re-expressed. **C.** Immunofluorescence images of MAD1 and MAD1-pT716 kinetochore levels in nocodazole-arrested HeLa FRT cells.

**Figure S6.**
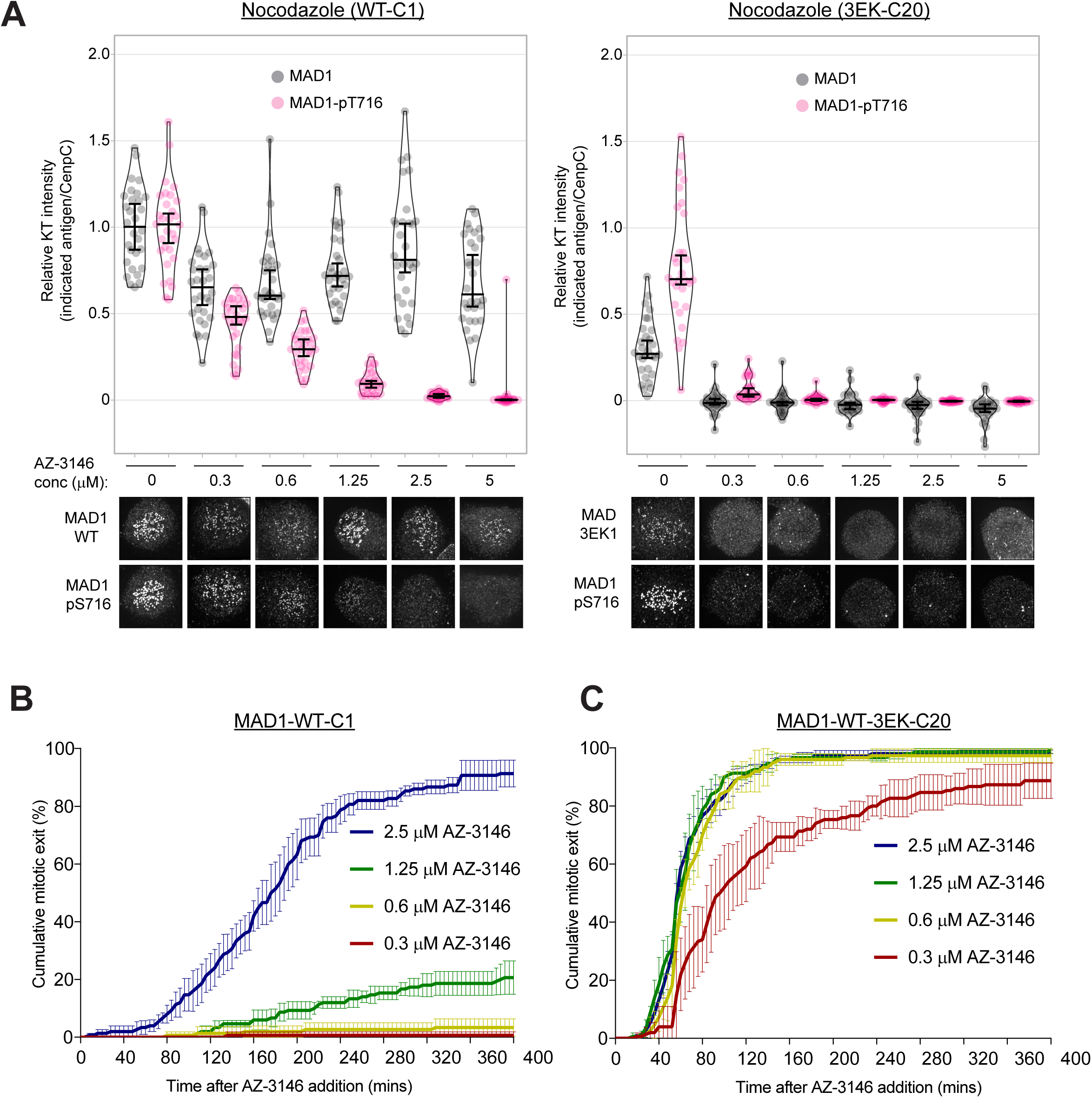
Corona MAD1 allows the SAC to tolerate MPS1 inhibition by preserving MAD1 at kinetochores and enhancing MAD1-pT716 levels. **A.** Quantifications (top) and corresponding immunofluorescence images (underneath) of kinetochore MAD1 and MAD1-pT716 levels in nocodazole-arrested MAD1-WT-C1 and 3EK-C20 treated with different doses of AZ-3146 for 30 min. MG132 was included at the time of AZ-3146 addition to prevent mitotic exit. Each dot represents a cell, horizontal lines indicate the median and error bars show 95% confidence interval. 30 cells from 3 experiments. **B**,**C.** Duration of mitotic arrest in MAD1-WT C1 (B) or MAD1-3EK C20 (C) cells arrested in nocodazole and then treated with indicated concentrations of AZ-3146. Graph shows cumulative mean (±SEM) of 3 experiments, 50 cells per condition per experiment. Some results are replicated in Figures 3D, 4D-E to allow a complete comparison of the full dose range.

